# Compensatory mutations reducing the fitness cost of plasmid carriage occur in plant rhizosphere communities

**DOI:** 10.1101/2022.08.01.502293

**Authors:** Susannah M. Bird, Samuel Ford, Catriona M. A. Thompson, Richard Little, James P.J. Hall, Robert W. Jackson, Jacob Malone, Ellie Harrison, Michael A. Brockhurst

## Abstract

Plasmids drive bacterial evolutionary innovation by transferring ecologically important functions between lineages, but acquiring a plasmid often comes at a fitness cost to the host cell. Compensatory mutations, which ameliorate the cost of plasmid carriage, promote plasmid maintenance in simplified laboratory media across diverse plasmid-host associations. Whether such compensatory evolution can occur in more complex communities inhabiting natural environmental niches where evolutionary paths may be more constrained is, however, unclear. Here we show a substantial fitness cost of carrying the large conjugative plasmid pQBR103 in *Pseudomonas fluorescens* in the plant rhizosphere. This plasmid fitness cost could be ameliorated by compensatory mutations affecting the chromosomal global regulatory system *gacA/gacS*, which arose rapidly in plant rhizosphere communities and were exclusive to plasmid carriers. These findings expand our understanding of the importance of compensatory evolution in plasmid dynamics beyond simplified lab media. Compensatory mutations contribute to plasmid survival in bacterial populations living within complex microbial communities in their natural environmental niche.

## Background

Plasmids play an important role in bacterial evolution by transferring ecological functions between lineages and thus driving genomic divergence [1-4]. Acquiring a new conjugative plasmid is, however, often costly for the host cell [5]. Such fitness costs are caused by a variety of mechanisms including genetic conflicts, regulatory disruption, cytotoxicity, codon use mismatches, or the metabolic burden of maintaining the plasmid [5-6]. Due to these fitness costs and the potential for segregational loss of the plasmid at cell division, plasmids are expected to be lost from bacterial populations inhabiting environments where plasmid-encoded functions are not beneficial [7]. Moreover, even in environments where plasmid-encoded functions are beneficial, these genes are expected to transfer to the chromosome, enabling loss of the plasmid [7]. Nonetheless, plasmids are common features of bacterial genomes, a situation termed the plasmid paradox [7–9].

Laboratory evolution experiments have shown that compensatory mutations that reduce the fitness costs associated with plasmid carriage can stabilize plasmids across diverse plasmid-host pairs (reviewed in [9]). Compensatory mutations can affect genes resident on the chromosome [10-15], the plasmid [14, 16, 17], or both [18-21] replicons. They target a wide range of gene functions across plasmid-host pairs, including regulators [11, 12, 22], RNA polymerase [12], helicases [15], other resident mobile genetic elements [23], or hypothetical genes without known functions [14]. Although commonly observed in simplified laboratory media, the importance of compensatory evolution for plasmid stability in more realistic, natural environments is far less well-studied.

In this study we ask whether compensatory mutations arise in response to plasmid carriage in bacterial communities living on plants. *Pseudomonas fluorescens* SBW25 is a plant commensal bacterium that was isolated from sugar beet in the 1990s at Wytham Woods, Oxfordshire, UK [24]. Around the same time, a large collection of conjugative mercury resistance plasmids was isolated by exogenous capture from the same field site [25], including the 425 kb conjugative megaplasmid pQBR103 [26]. SBW25 carrying pQBR103 can persist on plants both in the greenhouse and in the field [27], displaying an intriguing seasonal dynamic: plasmid carrying populations initially decline in abundance compared to plasmid-free populations, before recovering to roughly equal abundances later in the growing season (∼100 days after sowing).

The initial decline in plasmid-carrying SBW25 abundance relative to the plasmid-free population on plants [27] suggests that pQBR103 imposed a fitness cost. Subsequent experimental analysis showed pQBR103 acquisition imposes a large ∼50% fitness cost upon SBW25 both in nutrient broth and in potting soil [28], however, whether pQBR103 is costly on plants (i.e., SBW25’s and pQBR103’s natural ecological niche) is unknown. The recovery of plasmid-carrier abundance later in the growing season is also currently unexplained. The authors of the original study suggested that this dynamic may have been due to an unknown fitness benefit of the plasmid that is apparent only on mature plants [27]. An alternative hypothesis is that the recovery of plasmid carriers may have been driven by compensatory evolution to reduce the fitness cost of plasmid carriage. In nutrient broth, the cost of pQBR103 carriage is rapidly negated by compensatory mutations that target either the two component global regulatory system, *gacA/gacS*, or the gene of unknown function *PFLU4242*, which itself is positively regulated by *gacA/gacS* [11, 14]. Single mutations affecting either of these genes completely ameliorate the fitness cost of pQBR103 carriage in nutrient broth [14]. Whether these compensatory mutations arise or can ameliorate plasmid fitness costs on plants is unknown.

The GacA/GacS two component system works together with Rsm proteins to post-transcriptionally regulate hundreds of SBW25 chromosomal genes [29]. The SBW25 Gac-Rsm regulon includes a variety of traits that may affect colonisation and competitiveness on plants, including secreted secondary metabolites, motility, and biofilm formation[29], suggesting that loss of this system may be detrimental in complex natural environments such as plants. Interestingly, the pQBR103 sequence encodes an Rsm homologue, *PQBR443* (named *rsmQ*) [26, 28, 30]. RsmQ interacts with Gac-Rsm signalling within SBW25 cells to alter the expression of chemotaxis, motility, and metabolic phenotypes [30] and could thus alter the fitness effect of pQBR103 and the potential for compensatory mutation in plant rhizosphere communities.

To test if pQBR103 carriage imposes a fitness cost upon SBW25 living on plants and whether this cost can be ameliorated by known compensatory mutations, we competed plasmid carriers against plasmid-free SBW25 in the rhizosphere of wheat. Specifically, we tested the effect of plasmid carriage with or without *rsmQ* on competitive fitness of wild-type SBW25 and of SBW25 with a deletion of either the *gacS* gene or the *PFLU4242* gene. Next, we tracked the abundance of SBW25 with or without a plasmid for 4 weeks within bacterial communities inhabiting the rhizospheres of wheat plants. We quantified plasmid maintenance over time in SBW25 and used whole genome sequencing of clones isolated after 4 weeks on plants to determine if compensatory mutations had arisen in plasmid carrying bacteria. Our competition data showed that pQBR103 did indeed impose a large fitness cost on SBW25 in the wheat rhizosphere and that this cost was ameliorated by mutation of either of the known compensatory loci we tested. Both the fitness cost and the efficiency of amelioration by compensatory mutations were unaffected by the presence of *rsmQ*. Over 4 weeks, plasmid carriers typically reached lower abundance than plasmid-free SBW25 in the wheat rhizosphere community but stably maintained the plasmid at appreciable frequencies. Genome sequencing revealed that compensatory mutations affecting the *gacA/gacS* two component global regulatory system had arisen in multiple plasmid-carrying clones. Conversely, these genes were never mutated in plasmid-free controls. Further genome sequencing of earlier sampled clones revealed that compensatory mutations affecting *gacA/gacS* could be detected even after just 1 week on plants. These findings extend our understanding of the maintenance of costly plasmids from the lab into a more natural environment, showing that compensatory mutations rapidly and repeatedly occur in the plant rhizosphere. This suggests that evolution to compensate plasmid-imposed fitness costs may indeed promote the survival of plasmids in natural communities.

## Results

To quantify if acquisition of pQBR103 by SBW25 is costly on plants, and whether known compensatory mutations ameliorate fitness costs, we competed plasmid carriers against isogenic plasmid-free SBW25 in the wheat rhizosphere. We used 3 bacterial genetic backgrounds as plasmid carriers, wild-type SBW25 or mutants carrying deletions of either *gacS* or *PFLU4242*, enabling us to test the fitness effect of these known compensatory mutations on plants. In addition, we used two pQBR103 genotypes: wild-type pQBR103 or a mutant carrying a deletion of *rsmQ* (pQBR103ΔrsmQ). The inclusion of this mutant enables us to additionally test if interactions between RsmQ and the SBW25 Gac-Rsm system contributed to fitness costs on plants. Consistent with our previous experiments using nutrient broth or potting soil, we observed a large fitness cost of plasmid carriage in SBW25 in the wheat rhizosphere (figure 1; SBW25(pQBR103) vs SBW25: t = -4.868, p < 0.001; SBW25(pQBR103ΔrsmQ) vs SBW25: t = -4.883, p < 0.001). Moreover, the plasmid fitness cost in SBW25 was unaffected by deletion of *rsmQ* (pQBR103 vs pQBR103ΔrsmQ: t = -0.015, p = 1) and therefore subsequent analyses combined these plasmid treatments. These data show that pQBR103 does indeed cause a substantial fitness cost in SBW25 on plants, confirming that this plasmid is costly in an environment similar to that from which it was isolated (i.e., the plant rhizosphere). Deletion of either *gacS* or *PFLU4242* completely ameliorated the fitness cost of plasmid carriage (SBW25ΔgacS(pQBR103) vs SBW25: t = 0.808, p = 0.964; SBW25ΔPFLU4242(pQBR103) vs SBW25: t = 0.437, p = 0.998). These data show that both SBW25 compensatory mechanisms previously observed to evolve in nutrient broth function equivalently to negate fitness costs in the more natural environment of the plant rhizosphere. Intriguingly, although deletion of *PFLU4242* had no fitness effect in the absence of plasmids, deletion of *gacS* was beneficial in the absence of plasmids in competition with SBW25 (SBW25ΔgacS vs SBW25: t = 3.559, p = 0.0102). In line with previous studies of plant-associated *Pseudomonas* [31], this suggests that the Gac regulon is expressed by SBW25 on plants, that this expression is costly, but that non-responsive *gacS* mutants growing alongside wild-type cells may benefit from their Gac-regulated products.

**Figure 1.**
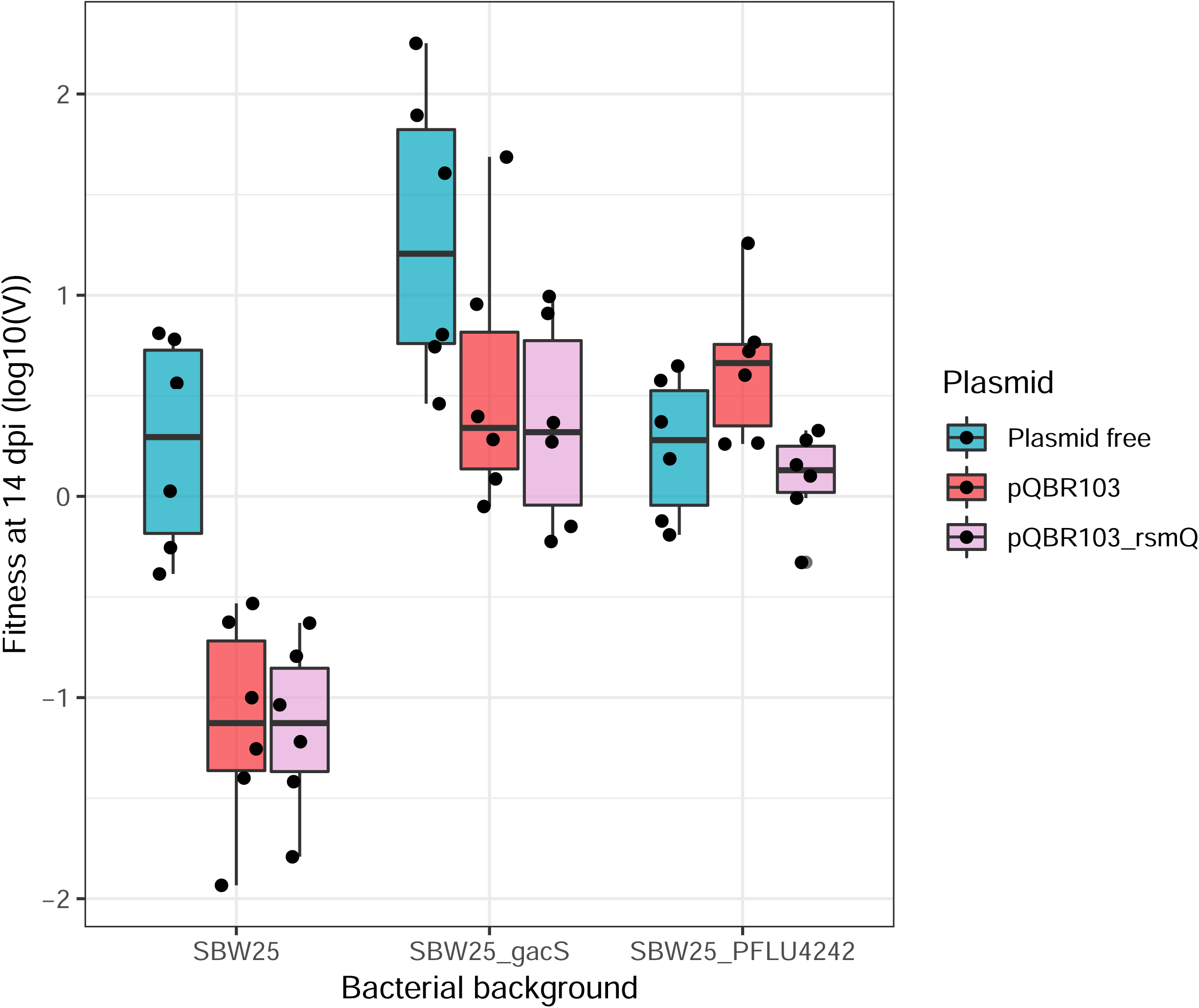
Fitness effects of pQBR103 carriage upon SBW25 with or without compensatory mutations in the wheat rhizosphere. Relative fitness (*v*) of strains from 3 bacterial backgrounds (SBW25, SBW25ΔgacS and SBW25ΔPFLU4242) either carrying no plasmid (blue), a wild type plasmid (red) or a *rsmQ* knockout plasmid (pink) was measured in direct competition with plasmid free wild type SBW25 in the wheat rhizosphere after 14 days. Boxplots show mean and interquartile range with replicate values shown as points in black (n=6).

By screening for transconjugants in the originally plasmid-free SBW25 competitor populations at 14 days, we found evidence for conjugation of both pQBR103 and pQBR103ΔrsmQ plasmids in the rhizosphere in all replicates. Transconjugant frequencies reached up to ∼1.5% of the originally plasmid free SBW25 competitor, and were highest from donors that carried compensatory mutations (figure S1, SBW25_gacS vs SBW25: t = 4.830, p < 0.001, SBW25_PFLU4242 vs SBW25: t = 8.579, p < 0.001) consistent with their higher fitness (t = 4.454, p < 0.0001, R^2^ = 0.349). There was no effect of plasmid genotypes on transconjugant frequency (F = 0.0251, p = 0.875). These data confirm that pQBR103 transfers by conjugation in the rhizosphere and further show that horizontal transmission of the plasmid is promoted by host compensatory mutations.

We next tested if plasmid carriage altered the ecological dynamics of SBW25 when in a bacterial community inhabiting the wheat rhizosphere. Wheat seedlings were colonized with SBW25, SBW25(pQBR103) or SBW25(pQBR103Δ*rsmQ*) alongside a background microbial community previously plated onto nutrient agar from root washes of mature wheat plants. Plants were destructively sampled weekly for 4 weeks and rhizosphere communities were plated onto selective agar to distinguish SBW25 populations from the background microbial community and plasmid carriers from plasmid-free cells. Plasmid carriage reduced the abundance of SBW25 in the wheat rhizosphere (figure 2A; SBW25(pQBR103) vs SBW25: t = 3.22, p = 0.005; SBW25(pQBR103Δ*rsmQ*) vs SBW25: t = 3.17, p = 0.006) with no effect of plasmid genotype (SBW25(pQBR103) vs SBW25(pQBR103Δ*rsmQ*): t = 0.005, p = 0.999). The presence of SBW25 with or without plasmids had no effect on the abundance of the background microbial community inhabiting the wheat rhizosphere (figure S2: F = 2.219, p = 0.0894). Both plasmids remained at appreciable frequencies in the SBW25 population for the duration of the experiment (figure 2B), although pQBR103 declined to lower prevalence than pQBR103Δ*rsmQ* during the later stages of the experiment (t = 4.061, p = 0.0001). These data suggest, consistent with our relative fitness estimates, that plasmid carriage is costly in the plant rhizosphere, but that plasmids are stably maintained nonetheless.

**Figure 2.**
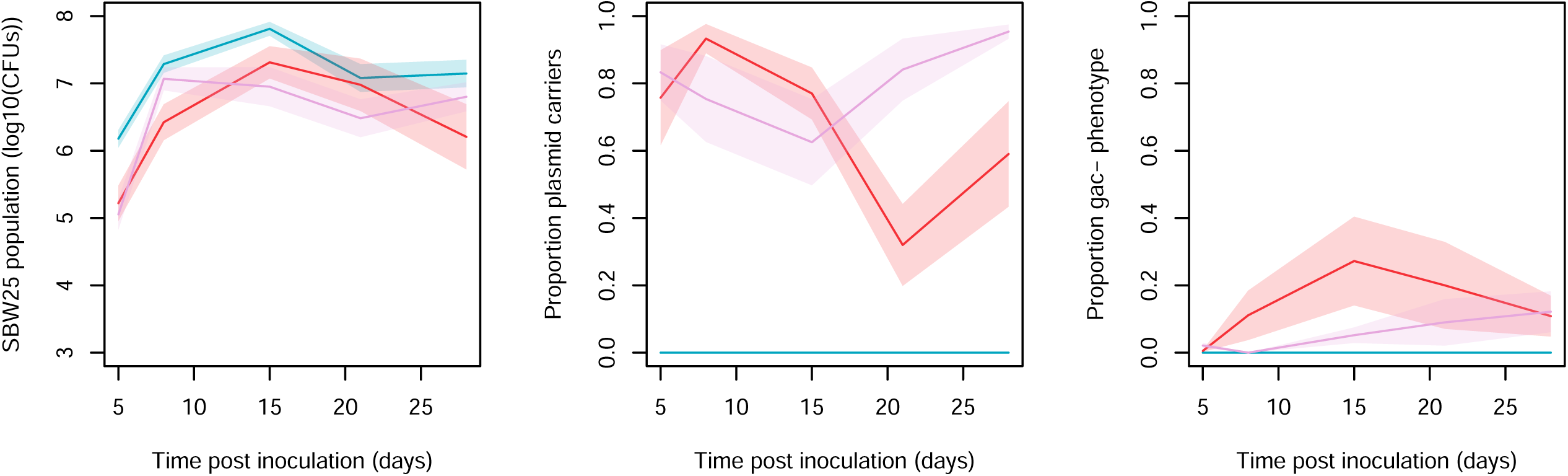
Dynamics of the SBW25 populations in the wheat rhizosphere over time. A. Total SBW25 population density. B. Proportion of SBW25 plasmid carriers within the population. C. Proportion of Gac-negative phenotypes in the SBW25 population. Lines denote means with standard error shown as the area around the mean. SBW25 populations were founded from clones carrying either no plasmid (blue), the wild type pQBR103 (red) or pQBR103ΔrsmQ (pink).

At the final sampling point, we picked one random SBW25 colony per plant for whole genome sequencing to determine the genetic response to selection and how this varied among treatments. We observed 11 mutations across the 22 sequenced genomes, with individual genomes containing between 0 and 2 mutations (Figure 3). All observed mutations affected chromosomal loci and no mutations were observed on plasmids. The only function subject to parallel mutations arising independently in multiple genomes, which is a signal of natural selection having acted upon these mutations, was *gacA/gacS* and these were exclusive to plasmid carrying clones. Specifically, *gacA* was mutated in one evolved clone carrying pQBR103 whilst *gacS* was mutated in two evolved clones carrying pQBR103ΔrsmQ (Figure 3). Because the SBW25-selective agar plates had been supplemented with skimmed milk powder we were able to detect the activity of a secreted protease that is positively regulated by *gacA/gacS*, enabling us to track the emergence of phenotypically Gac-negative SBW25 mutants over time [32]. These data reveal that Gac-negative SBW25 colonies were detected only in the presence of plasmids, but never in their absence, and arose rapidly (within 1-2 weeks; Figure 2) (effect of time; t = 5.113, p < 0.001) reaching intermediate frequencies in SBW25 populations containing either plasmid genotype. Colonies with a Gac-negative phenotype were marginally more common in populations containing pQBR103 compared to pQBR103ΔrsmQ (t = -2.159, p = 0.0349; Figure 2). Genome sequencing of some of these plasmid-carrying phenotypically Gac-negative SBW25 colonies confirmed that the majority (8 out of 9 colonies) contained mutations affecting either *gacA* (n=2) or *gacS* (n=6) (Figure S3). These data show that mutation of *gacA/gacS* was exclusively associated with plasmid carriage and suggest, in combination with our relative fitness data, that such mutations arose to compensate for the fitness cost of carrying the plasmid in the plant rhizosphere.

**Figure 3.**
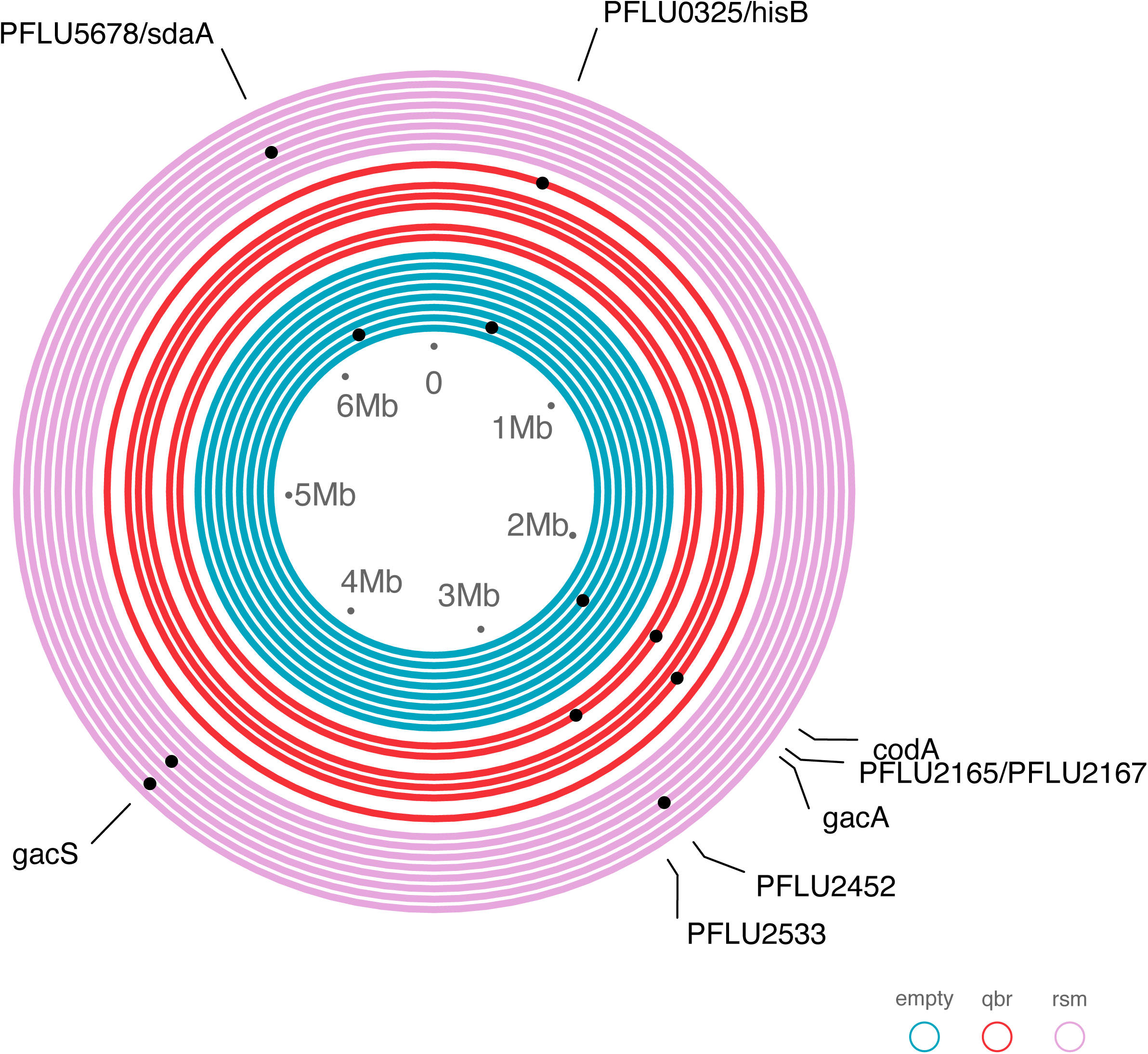
Genome plots showing the chromosomal mutations observed in evolved clones sampled on day 28 of the plant experiment. Concentric rings represent the chromosome of individual sequenced clones colour-coded by treatment - SBW25 (blue), SBW25(pQBR103) (red) and SBW25(pQBR103Δ*rsmQ)* (pink). Black dots denote the location of mutations with gene targets shown on the outer labels.

## Discussion

Laboratory evolution experiments suggest that compensatory evolution to reduce the cost of plasmid carriage plays an important role enabling the persistence of plasmids in bacterial populations [9]. However, whether similar evolutionary processes operate in more complex natural settings, such as plant rhizosphere communities, is poorly understood. Indeed, the genomic flexibility afforded by rich lab media might be expected to be far more constrained in natural environments, potentially limiting the accessibility of compensatory mutations. Here we show, using the plant commensal bacterium *P. fluorescens* SBW25 and the environmental mercury resistance plasmid pQBR103, that although plasmid carriage is costly in the plant rhizosphere, such fitness costs can be ameliorated by compensatory mutations. Specifically, mutations affecting *gacA/gacS* arose rapidly and exclusively in plasmid carrying SBW25 populations, where they completely ameliorated the fitness cost of carrying pQBR103 in the plant rhizosphere.

Using competition experiments with defined mutants we show that loss of either the two component system *gacA/gacS* or the hypothetical gene *PFLU4242* is sufficient to reduce the fitness cost of carrying pQBR103 on plants. Our previous studies have shown that the pQBR103 fitness cost arises from a specific genetic conflict between *PFLU4242* and genes on pQBR103, inducing upregulation of chromosomal mobile genetic elements including prophages that are thought to cause cell damage [14]. Surprisingly, we did not observe mutation of *PFLU4242* in evolved plasmid carrying clones, despite mutation of this gene offering complete amelioration on plants. Instead, *de novo* compensatory mutations targeted *gacA/gacS*. Loss of the *gacA/gacS* two component system likely reduces plasmid fitness cost by downregulating *PFLU4242*, providing an indirect regulatory route to amelioration [14]. The predominance of *gacA/gacS* mutations is consistent with *in vitro* studies using this system where, although *PFLU4242* mutations were observed, they were much less common than mutations affecting *gacA/gacS* [11, 14]. It is possible that *gacA/gacS* merely represents a larger mutational target than PFLU4242, making mutations at these loci more likely to arise. Similarly, it has been argued that *gacA/gacS* may represent contingency loci [33–35] with elevated mutation rates [33, 34], and thus would be predicted to acquire mutations more frequently than other regions of the genome.

Gac mutants have been reported in natural *Pseudomonas* populations isolated from both soil and plants (reviewed in [34]), where they may gain a fitness benefit from not paying the metabolic costs of expressing Gac regulated genes, whilst benefiting from the secreted metabolites produced by neighbouring wild-type cells[36, 37] (although cf. [38] for an alternative view). Indeed, our competition experiments suggested that in addition to ameliorating the fitness cost of pQBR103, loss of *gacA/gacS* was also beneficial in the absence of plasmids. However, *de novo* Gac-negative mutants were only observed in plasmid carriers, and never in plasmid-free SBW25 populations in the plant rhizosphere. As such, Gac mutations appear to have been insufficiently beneficial to drive loss of Gac signalling by SBW25 in rhizosphere communities in the absence of plasmids. Compensatory evolution in response to plasmid carriage may therefore help to explain the variability of Gac phenotypes observed within rhizosphere communities [34, 38].

Although the plasmid genotypes did not differ in their fitness costs, we observed subtle differences in their dynamics and compensatory evolution. pQBR103ΔrsmQ was maintained at higher frequency in SBW25 over time compared to pQBR103, whereas Gac-negative mutants were marginally more common in pQBR103 than pQBR103ΔrsmQ containing SBW25 populations. Moreover, whereas a mixture of *gacA* and *gacS* mutations occurred in pQBR103 carriers, we only ever observed *gacS* mutations in pQBR103ΔrsmQ carriers. These results are consistent with our earlier findings that RsmQ interferes with the host Rsm pathway [30], which is itself induced by *gac* loss-of function mutations. Furthermore, *rsmQ* deletion recovers certain plasmid-induced phenotypes in SBW25 [30], which may slightly reduce the fitness cost of plasmid carriage in some environments.

Our data suggest an explanation for the previously reported seasonal dynamics of SBW25(pQBR103) on plants in the field, wherein plasmid carriers declined after sowing before rebounding later in the growing season on sugar beets [27]. The large competitive fitness cost of the plasmid that we measure in the rhizosphere is consistent with the reduced abundance of plasmid carriers compared to plasmid-free SBW25 we observed in rhizosphere communities, and as such this fitness cost seems the most likely reason for the initial decline of SBW25(pQBR103) observed by Lilley and Bailey on plants in the field. Compensatory mutations that completely ameliorate the fitness cost of the plasmid arise and spread to appreciable frequencies within weeks in plasmid carrier populations of SBW25 within rhizosphere communities. Such large fitness effects and rapid timescales of compensatory mutations in the rhizosphere is consistent with the reinvasion of SBW25(pQBR103) in the field after 100 days being driven, at least in part, by compensatory evolution. Although we cannot rule out the contribution of other plasmid genes providing a delayed benefit of pQBR103 later in the growing season as proposed in the original study [27], the dynamics of compensatory evolution we report here strongly suggest that such unknown fitness benefits are not necessary to explain the field dynamics.

This study expands our understanding of the importance of compensatory evolution in plasmid dynamics from simplified lab media to a more complex and plant-associated natural environment. Rapid compensatory evolution to reduce the fitness costs of plasmid carriage is likely to enable the stable persistence of costly plasmids in plant rhizosphere communities as well as in a range of other natural microbial communities [39].

## Materials and methods

### Plant varieties and culture conditions

Wheat plants (Skyfall variety, RATG) were grown in 50ml Falcon tubes containing 40cm^3^ of washed autoclaved vermiculite. 10 ml 1x Jensens nutrient solution was added to each microcosm and re-autoclaved. Wheat seeds were sterilised by agitating 5g in 30ml 30% bleach/0.01% TritronX solution for 10 minutes at room temperature. Bleach solution was removed by washing ten times in 50ml sterile water. Seeds were stratified overnight at 4℃and then spread onto sterile filter paper to germinate in the dark for 48 hours. Single seedlings were placed into sterile vermiculite microcosms and allowed to establish for 48 hrs in sterile conditions (lid on tube) prior to inoculation with bacterial populations/communities. Plants were grown in growth chambers (Conviron) at 250 *μ*mol m^-2^ s^-1^, 16:8 hrs light:dark cycle, 22℃day/18℃night and 60% relative humidity.

### Bacterial strains, plasmids, and culture conditions

The bacterial strains and plasmids used in this study are given in table 1. Plasmids were introduced into bacterial strains by conjugation and confirmed by PCR as previously described[28]. Overnight cultures were grown in 6 ml KB liquid medium in 30 ml glass universals shaken at 28 °C. Colony counts were performed on selective KB agar plates supplemented with Streptomycin (250 *μ*g/ml) or Gentamicin (30 *μ*g/ml) plus 50 *μ*g/ml X-gal to distinguish Sm^R^lacZ and Gm^R^ labelled *P. fluorescens* strains or with Kanamycin (50 *μ*g/ml) to select for plasmid carriers. Plates were incubated at 28 ℃for 48 hrs. To detect protease producing *P. fluorescens* colonies - i.e. those with a functional *gacAS* system -KB agar was supplemented with skimmed milk powder (250 *μ*g/ml) [32].

**Table 1.** Bacterial strains and plasmids.

| Bacterial strain or plasmid | Reference |
| --- | --- |
| *Pseudomonas fluorescens* SBW25 | [24] |
| *P. fluorescens* SBW25-Gm<sup>R</sup> | [28] |
| *P. fluorescens* SBW25-Sm<sup>R</sup>lacZ | [28] |
| *P. fluorescens* SBW25-Gm<sup>R</sup>Δ*gacS* | [11] |
| *P. fluorescens* SBW25-Gm<sup>R</sup>Δ*PFLU4242* | [12] |
| pQBR103 | [25] |
| pQBR103-Kan<sup>R</sup> | [13] |
| pQBR103-Kan<sup>R</sup>Δ*rsmQ* | [30] |

A standardised wheat rhizosphere community inoculum was generated from 6 week old wheat plants. Plants were grown in 2 litre pots containing John Innes Number 2 compost (Arthur Bower) for 6 weeks in a Conviron growth chamber at 250 *μ*mol m^-2^ s^-1^, 16:8 hrs light:dark cycle, 22℃day/18℃night and 60% relative humidity. The roots were destructively harvested and washed in 100mM Tris-phosphate buffer (pH 5.4) plus 1.8 mg/ml lipase (Sigma), 100 *μ*g/ml β-Galactosidase (Merck) and 4 *μ*g/ml α-Glucosidase (Merck) and incubated at 37℃for 30 mins, then vortexed with glass beads for 5 mins at room temperature. The root wash was diluted with an equal volume of sterile water, mixed and allowed to settle on ice for 10 mins. The wash was then centrifuged at 1000g for 30 secs and the supernatant carefully removed. The total bacterial rhizosphere community was quantified by plating out serial dilutions onto 0.1x NB agar plates supplemented with cycloheximide (1 mg/ml) to suppress fungal growth and 50 *μ*g/ml X-gal and incubated at 28 ℃for 48 hrs. Frozen aliquots of the rhizosphere community were stored in 10% glycerol at -80 ℃for subsequent use in plant rhizosphere dynamics experiments.

### Plant rhizosphere competition assays

Overnight KB cultures of the relevant *P. fluorescens* strains were subcultured into M9 pyruvate minimal liquid medium and pre-conditioned overnight shaken at 28 °C. These cultures were then centrifuged for 10 mins at 3500 g at 4℃and cell pellets resuspended in M9 salts solution. Competing strains were mixed at 1:1 ratios and plated onto X-gal to distinguish between strains. Six replicate gnotobiotic wheat seedling microcosms were inoculated per competition with 1 ml of mixtured culture under sterile conditions (total inoculated bacterial cells = approx. 1×10^4^ colony forming units). Plants were propagated in a Conviron growth chamber and destructively harvested after 14 days. To recover the bacterial populations, plant roots were separated from aerial tissue and transferred to Falcon tubes containing 10 ml of sterile 10mM MgSO4 0.1% Tween (root wash solution) and 2 cm^3^ of sterile glass beads. The roots were vortexed for 3minutes at room temperature, left for 30 minutes on ice and then vortexed again for 3 minutes. The root wash supernatant was removed and serial dilutions plated onto KB agar plates supplemented with 50 *μ*g/ml X-gal to distinguish the competing strains and incubated for 48 hr at 28°C. The relative fitness of a focal strain was calculated as *v* given by *v =* x_2_ (1 − x_1_) / x_1_ (1 − x_2_), where x_1_ is the starting proportion of the focal strain and x_2_ is the final proportion of the focal strain [40].

### Plant rhizosphere dynamics experiment

Seedling microcosms were inoculated with 1ml 6×10^6^ CFU/ml of either SBW25, SBW25(pQBR103), SBW25(pQBR103Δ*rsmQ*) or an equal volume of sterile water and allowed to establish for 5 days. Plants were then inoculated with approximately 10.5×10^6^ colony forming units of the standardised rhizosphere community. On days 5, 8, 15, 21, and 29, 6-8 plant microcosms were destructively harvested per treatment. Roots were processed as described above. Serial dilutions were plated onto selective agar plates to enumerate the total bacterial community (0.1x NB agar), the total *P. fluorescens* population and the frequency of protease producers (KB+streptomycin+skimmed milk), and the frequency of plasmid carriers (KB+streptomycin+kanamycin). Agar plates were incubated at 28℃for 48 hours prior to colony forming unit counts.

### Genomic analysis

One random *P. fluorescens* colony per plant sampled on day 29 and several colonies of protease-negative *P. fluorescens* sampled from plants across the experiment were chosen for whole genome sequencing. Whole-genome sequencing was performed by MicrobesNG using a 250bp paired-end protocol on the Illumina HiSeq platform. Variants were called against the ancestral reference genome using the Breseq computational pipeline using standard default settings [41]. All variants were validated visually using the alignment viewer IGV [42]. All sequencing data are available on the Sequencing Read Archive under accession PRJNA839377.

### Statistical analysis

All analysis was performed in R statistical software. Competition data was analysed using a linear model (lm(), base R) and further interrogated using Tukey Test multiple comparisons (glht(), multcomp). As comparisons showed no effect of the plasmid type on fitness a further analysis was performed grouping plasmid treatments to reduce the number of multiple comparisons. For the plant rhizosphere dynamics data, whole community and SBW25 population density were analysed using a linear model (lm(), base R) and where appropriate treatments were compared using Tukey’s pairwise comparisons (lsmeans(), library{emmeans}) Proportion data was analysed from counts using generalised linear model (glm(), base R) with a quasi binomial variance structure.

## Data Access Statement

All experimental data sets are provided in the Supplementary Information of this article. All genome sequence data is freely available from the Sequence Read Archive accession PRJNA839377.

## Figure Legends

**Figure S1.**
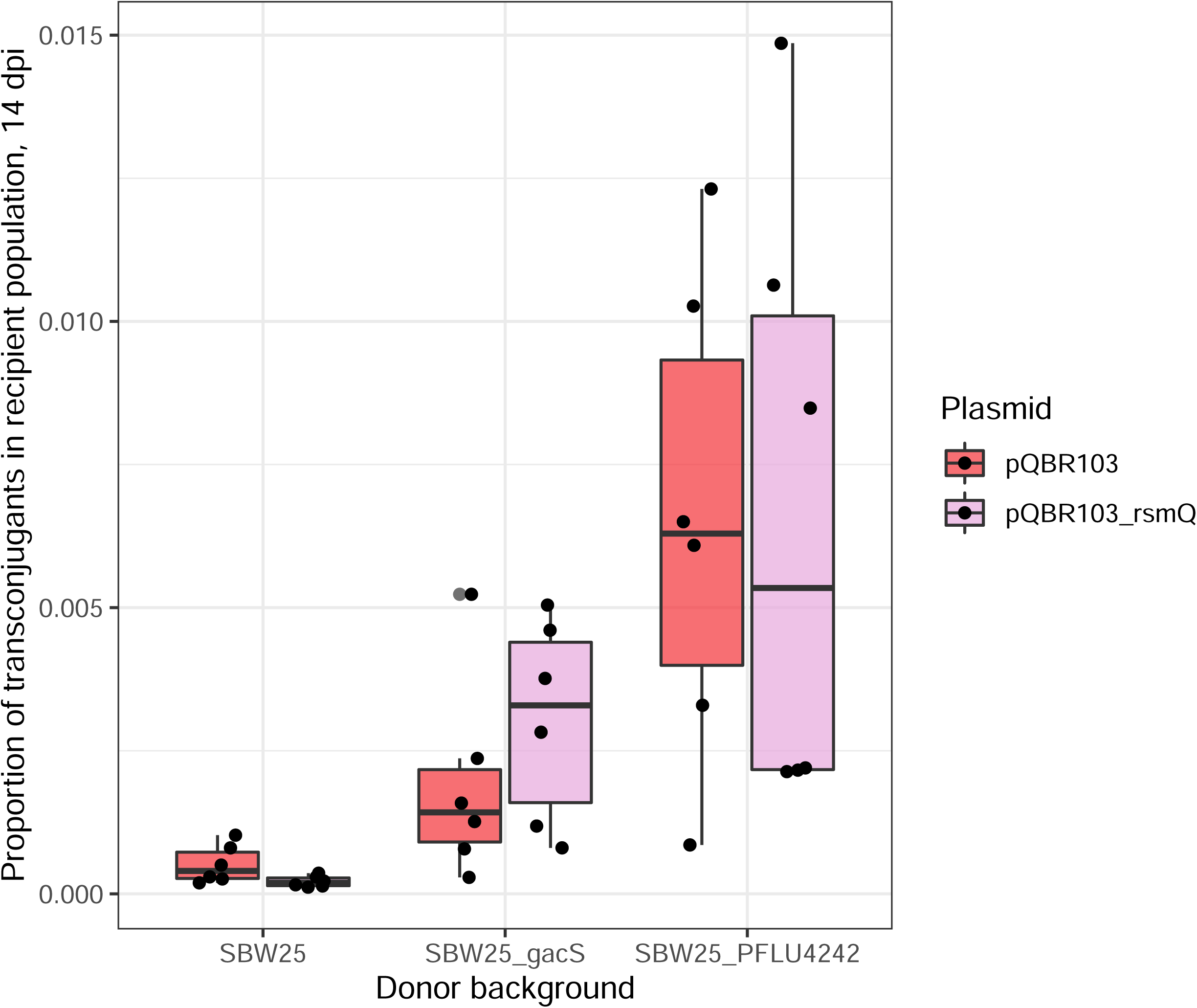
The proportion of transconjugants in initially plasmid free SBW25 competitor population after 14 days post infection. Plasmid carrying donor strains were either the wild type SBW25, SBW25ΔgacS or SBW25ΔPFLU4242. Strains carried either a wild type plasmid (red) or a rsmQ knockout plasmid (pink). Boxplots show mean and interquartile range with replicate values shown as points in black (n=6).

**Figure S2.**
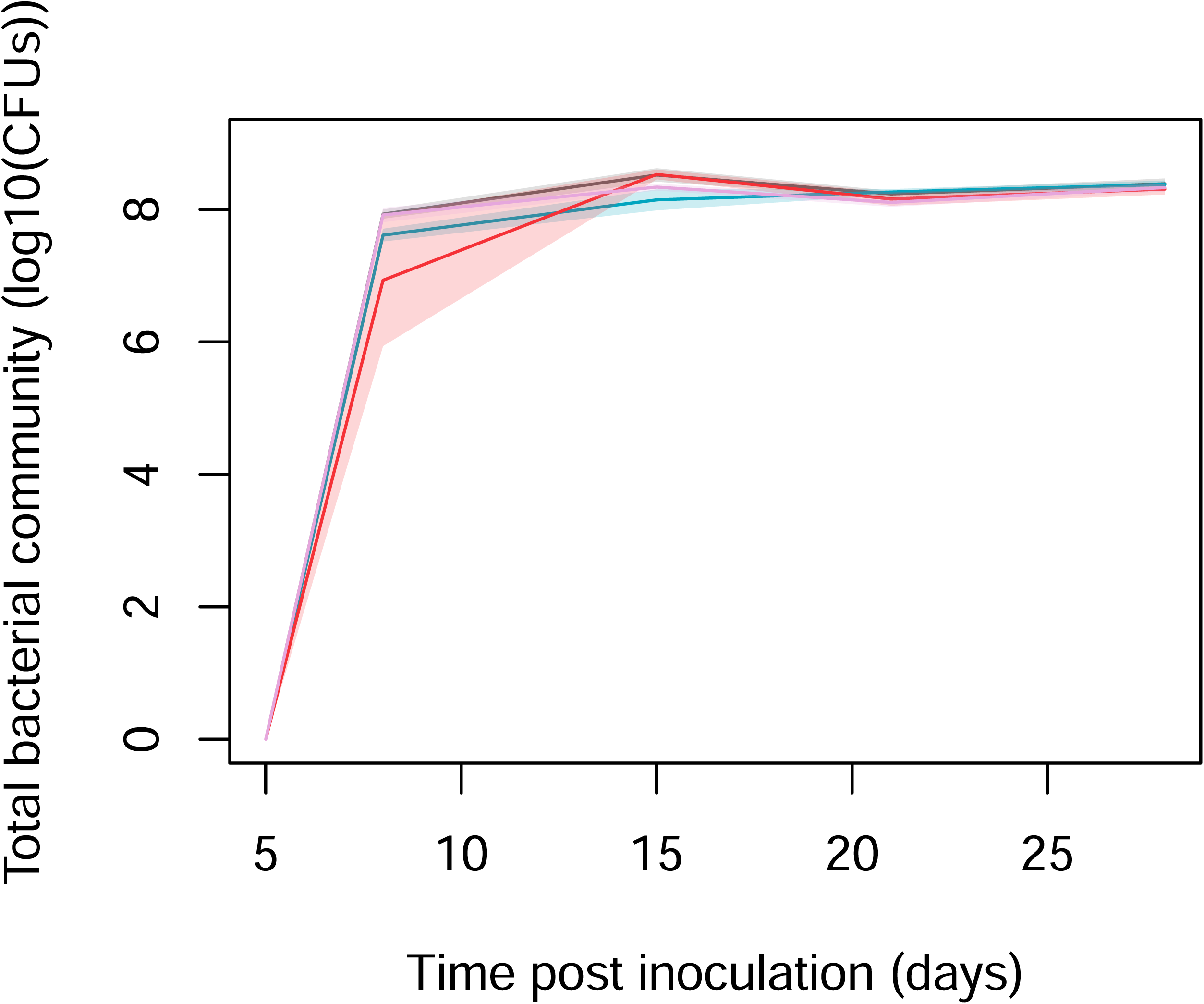
Dynamics of bacterial background community in the wheat rhizosphere. The density of non-SBW25 population was estimated by plating root wash on Xgal-agar. Because the SBW25 strain carries a *lacZ* gene then colonies that are not blue were assumed to be non-SBW25 belonging to the background community. Lines show means for 4 treatments; community only control (grey), SBW25 containing (blue), SBW25(pQBR103) containing (red) and SBW25(pQBR103ΔrsmQ) containing (pink). The shaded area denotes standard error.

**Figure S3.**
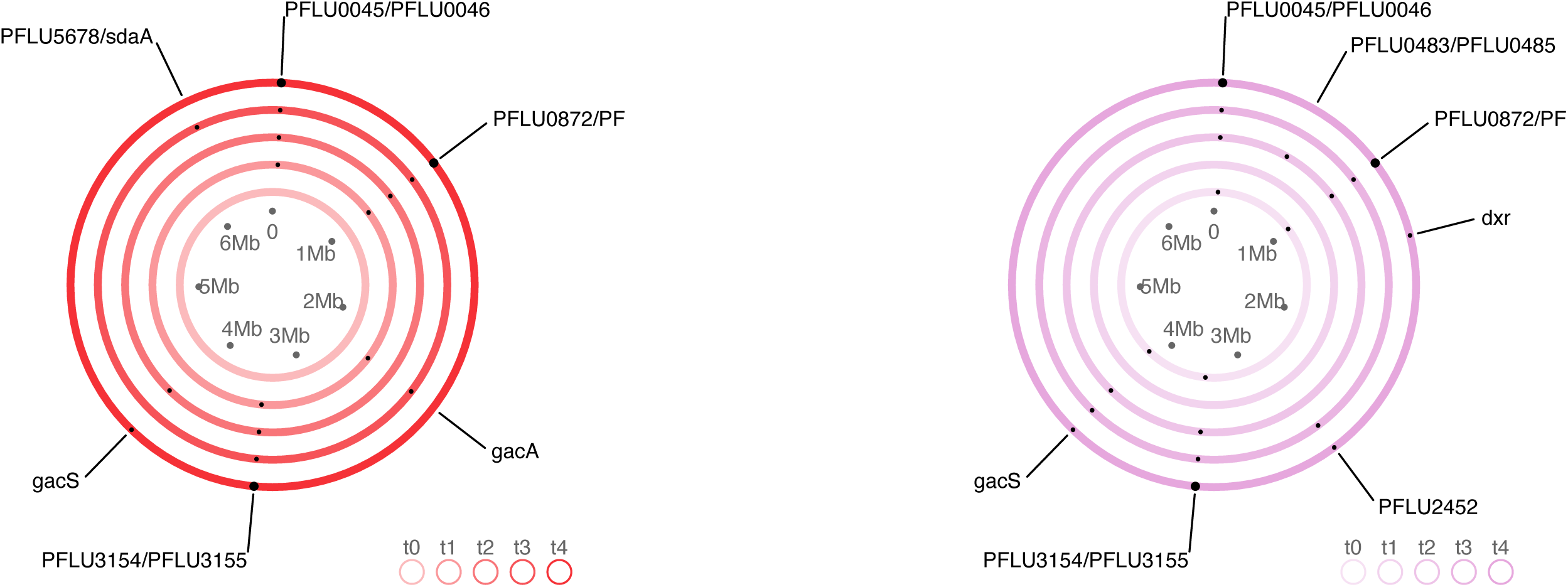
Genome plots for Gac-negative phenotype SBW25 evolved clones sampled over time. Concentric rings represent the chromosome of individual sequenced clones colour-coded by treatment, SBW25(pQBR103) (red) and SBW25(pQBR103Δ*rsmQ)* (pink). The sampling day when the clone was picked is denoted by shading. Black dots denote the location of mutations with gene targets shown on the outer labels.

